# DeMixSC: a deconvolution framework that uses single-cell sequencing plus a small benchmark dataset for improved analysis of cell-type ratios in complex tissue samples

**DOI:** 10.1101/2023.10.10.561733

**Authors:** Shuai Guo, Xiaoqian Liu, Xuesen Cheng, Yujie Jiang, Shuangxi Ji, Qingnan Liang, Andrew Koval, Yumei Li, Leah A. Owen, Ivana K. Kim, Ana Aparicio, John Paul Shen, Scott Kopetz, John N. Weinstein, Margaret M. DeAngelis, Rui Chen, Wenyi Wang

## Abstract

Bulk deconvolution with single-cell/nucleus RNA-seq data is critical for understanding heterogeneity in complex biological samples, yet the technological discrepancy across sequencing platforms limits deconvolution accuracy. To address this, we introduce an experimental design to match inter-platform biological signals, hence revealing the technological discrepancy, and then develop a deconvolution framework called DeMixSC using the better-matched, i.e., benchmark, data. Built upon a novel weighted nonnegative least-squares framework, DeMixSC identifies and adjusts genes with high technological discrepancy and aligns the benchmark data with large patient cohorts of matched-tissue-type for large-scale deconvolution. Our results using a benchmark dataset of healthy retinas suggest much-improved deconvolution accuracy. Further analysis of a cohort of 453 patients with age-related macular degeneration supports the broad applicability of DeMixSC. Our findings reveal the impact of technological discrepancy on deconvolution performance and underscore the importance of a well-matched dataset to resolve this challenge. The developed DeMixSC framework is generally applicable for deconvolving large cohorts of disease tissues, and potentially cancer.

## Introduction

Although recent advances in single-cell/nucleus RNA sequencing (sc/snRNA-seq) offer valuable insights into cell types and states in healthy^1^ and diseased tissues^2,3^, high expense and complex sample preparation procedures have restricted its widespread adoption in clinical settings^4^. Bulk RNA-seq, on the other hand, retains its essential role, especially in large disease-based cohort studies, for which its cost-efficiency, streamlined sample processing, and high-throughput analytic capabilities establish it as the method of choice for both preliminary screenings and exhaustive population-level analyses^5–7^. Nevertheless, bulk RNA-seq comes with a significant drawback: it captures averaged gene expression across heterogeneous cell types, thus confounding downstream analysis^4^. To mitigate this drawback, deconvolution methods have been developed to delineate the cell-type-specific signals from bulk RNA-seq data. Traditional bulk-based deconvolution methods^8,9^ employ bulk RNA-seq data from normal tissues or cell lines as the reference. They were typically constrained by low-resolution estimates, limited to identifying only two or three cellular components within the bulk samples. The progress of sc/snRNA-seq techniques opens the door to the emergence of single-cell-based deconvolution methods^10–18^, which tap into the granularity of even a modest set of single-cell data to provide far superior resolution in estimating cell-type proportions in complex tissues, thereby offering a cost-effective alternative.

Single-cell-based deconvolution methods are not without their disadvantages, however. Though affording remarkable resolution, they encounter a substantial challenge in achieving precision and accuracy. The limitations arise from inconsistencies in gene expression profiles between bulk and sc/snRNA-seq data. Those inconsistencies are attributable to technique variations in sample acquisition, preparation, and sequencing^19–22^. Such inconsistencies, which we refer to as “technological discrepancies”, have caused prior deconvolution studies to produce suboptimal estimates of cell-type proportions, particularly when unpaired sc/snRNA-seq data serve as the reference matrix for deconvolving publicly available large bulk cohorts^16,23,24^. Researchers have become aware of these discrepancies^25–27^, and several studies have attempted to address these issues but ended up with limited success^10,14,18^. CIBERSORTx^18^ simply implements a batch effect correction step but offers limited improvements in deconvolving complex bulk tissues. The ensemble approach of SCDC^14^ uses matched bulk and scRNA-seq data from two normal tissue samples (e.g., mouse breast) but lacks generalizability to patient cohorts. The most recent SQUID^10^ builds on top of DWLS^15^ with a Bisque-based linear transformation step^28^ to align matched bulk and scRNA-seq data; it can distort gene expression profiles, risking overcorrection. Meanwhile, existing benchmarking designs^10,23,24,29^ often employ datasets such as simulated pseudo-bulk data, cell line mixtures, or publicly available data, none of which are tailored to reveal the negative effect of technological discrepancy. Therefore, there is an unmet need for a comprehensive benchmark study that rigorously illustrates and assesses the impact of technological discrepancy and a robust deconvolution framework that effectively mitigates such discrepancy to enhance analytical accuracy.

In this paper, we thoroughly explore the technological discrepancy and address its impact on deconvolution performance to make adjustments correspondingly. To accomplish this, we generate a specialized benchmark dataset of 24 healthy retinal samples, ensuring technological discrepancy as the main confounding factor. Using this dataset, we demonstrate that technological discrepancy significantly affects the expression profiles of bulk and single-nucleus data and thus reduces the accuracy of existing single-cell-based deconvolution methods. Against this backdrop, we introduce a novel deconvolution method called DeMixSC, which employs a benchmark dataset and an improved weighted nonnegative least-squares (wNNLS) framework^30^ to identify and adjust for genes consistently affected by technological discrepancy. DeMixSC is generalizable to any tissue type, given a small representative benchmark dataset, to effectively deconvolve a large tissue-type-matched bulk cohort. We validated the improved deconvolution performance of DeMixSC by comparing it on our benchmark dataset with eight other existing methods. When applied to 453 peripheral retinal samples from patients with age-related macular degeneration (AMD)^7^, DeMixSC achieved more realistic cell-type estimates that reflect subtle changes in cell-type proportions among AMD grades, indicating that it can be reliable and generalizable in real-world settings. DeMixSC fills the gap in resolving the technological discrepancy in bulk deconvolution and serves as an accurate and adaptable tool for estimating cell-type proportions.

## Results

### Use benchmark data to assess technological discrepancy

We designed and generated a specialized benchmark dataset to assess the technological discrepancy between bulk and sc/sn sequencing platforms (Fig. 1A). This dataset comprises 24 healthy retinal samples from donors’ eyes collected within six hours postmortem (ages of death between 53 and 91, Supplementary Table 1), for two batches of sequencing experiments. Both bulk and snRNA-seq profiling were performed on each sample from the same single-nucleus suspension aliquot using a template-switching method to generate full-length cDNA libraries (see Methods). Because single-cell protocols can be biased toward retaining certain cell types^31^, hence changing the cell-type proportions, this special approach maximizes our chance that the matched sequencing data share approximately the same cell-type proportions. We performed the cell-type annotation for snRNA-seq data with known markers (see Methods and Supplementary Table 2). The resulting snRNA-seq data is summed to create matched pseudo-bulk RNA-seq data (see Methods). We hypothesize that any major differences in gene expression profile between the matched pseudo-bulk and real bulk RNA-seq would be technological, rather than due to biological discrepancies.

**Figure 1.**
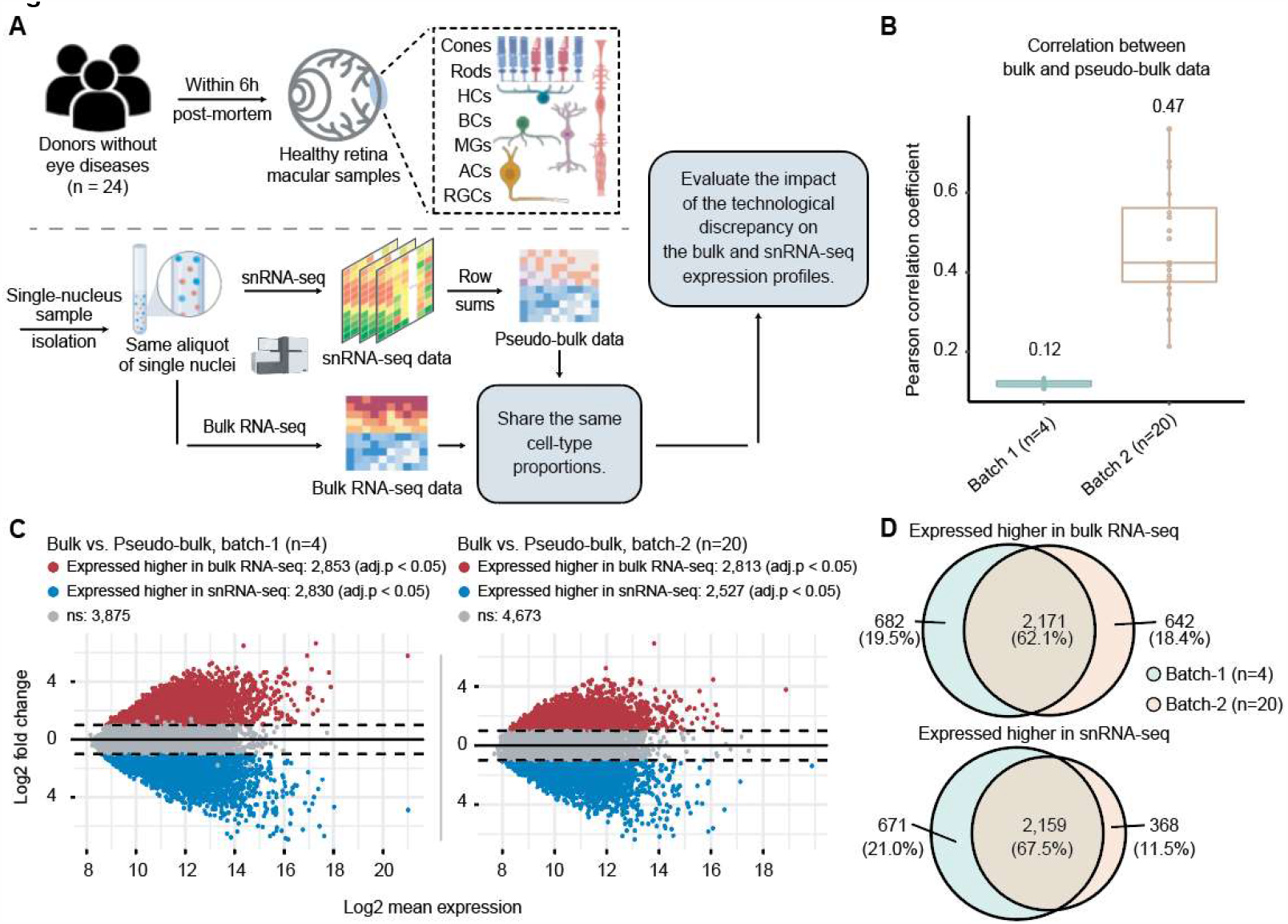
Assessing technological discrepancy between bulk and single-cell sequencing platforms using matched single-nuclei aliquots. **A**, Workflow for generating a benchmark dataset. We collect 24 healthy human retinal samples within six hours of postmortem. An illustration shows the layer and cell compositions of the human retina. Seven major cell types include photoreceptors (Rod and Cone cells), bipolar cells (BCs), retinal ganglion cells (RGCs), horizontal cells (HCs), amacrine cells (ACs), and Müller glia cells (MGs). Three minor cell types are not depicted in the illustration: astrocytes, microglia cells, and retinal pigment epithelial cells (RPEs). Samples are isolated into single-nucleus suspensions. The same aliquot of single-nuclei is used for both bulk and snRNA-seq profiling. The matched pseudo-bulk mixtures are generated as conventionally done by summing UMI counts across cells from all cell types in each sample. This data generation pipeline guarantees the matched bulk and snRNA-seq data share the same cell-type proportions, which enables us to evaluate the impact of technological discrepancy (i.e., the shot-gun sequencing procedure) on the bulk and snRNA-seq expression profiles. **B** and **C** show the influence of technological discrepancy at the sample- and gene-level, respectively. **B**, Pearson correlation coefficient across genes between the matched real-bulk and pseudo-bulk RNA-seq data for one sample at a time for both batches. **C**, MA-plots displaying the mean expression levels of all genes between matched real-bulk and pseudo-bulk data. Differentially expressed (DE) genes are identified using the paired t-test with Benjamini-Hochberg (BH) adjustment. Red represents genes expressed higher in the real-bulk, and blue represents genes expressed higher in the pseudo-bulk. The horizontal dotted lines denote a 2-fold change between matched real-bulk and pseudo-bulk data. adj.p: adjusted *P*-values. **D**, Venn diagrams showing genes consistently expressed higher in the bulk (upper) or the pseudo-bulk (bottom) between the two batches, which were generated using different tissue samples and at a different time.

We indeed observe much larger batch differences between real-bulk and pseudo-bulk data, than the small differences in cell-type distributions across samples in snRNA-seq or differences between the two experimental batches (Extended Data Figs. 1A-E). Total read counts from bulk RNA-seq data are significantly lower than total UMI counts from matched pseudo-bulk data (Extended Data Fig. 1F). Assuming that the difference in read depth does not impact the relative expression for each gene, we expect gene expression correlation to be a better metric for identifying technological discrepancy. We observe a low-to-moderate correlation of gene expression, consistent across samples, between the paired bulk data sets (Fig. 1B, mean Pearson correlation coefficient = 0.12 for batch-1, and 0.47 for batch-2). Further differential expression (DE) analysis between the two bulk samples identifies more than 5,000 DE genes in each experimental batch (Fig. 1C, adjusted *P*-values < 0.05), with more than 60% of those genes overlapping across the experiments (Fig. 1D). Our observations suggest a consistent technological effect across experiments. In broader contexts, factors such as library preparation, RNA capture efficiency, reverse transcription protocol, and sequencing depth could serve as potential sources of technological discrepancy^19,20,32^. We, therefore, expect that the reference matrices derived from sc/snRNA-seq data will not fully represent cell-type-specific expression profiles in bulk samples^10,24^. Given such discrepancies, the performance of existing deconvolution methods will be compromised, as their key assumption about the representative reference is violated.

### Overview of DeMixSC

Here, we present our novel deconvolution framework, DeMixSC, and illustrate how it addresses the observed consistent technological discrepancy in order to enhance the estimation accuracy of cell-type proportions. The DeMixSC framework, as depicted in Fig. 2, is built upon the commonly used wNNLS approach^15,17,30^ with several essential improvements (see Methods and Supplementary Note 1). Concretely, for a subject *j*, DeMixSC estimates its cell-type proportions, denoted by 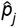, by minimizing a composite of two weighted squared error terms, as given in (1):

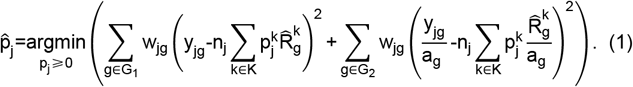

**Figure 2.**
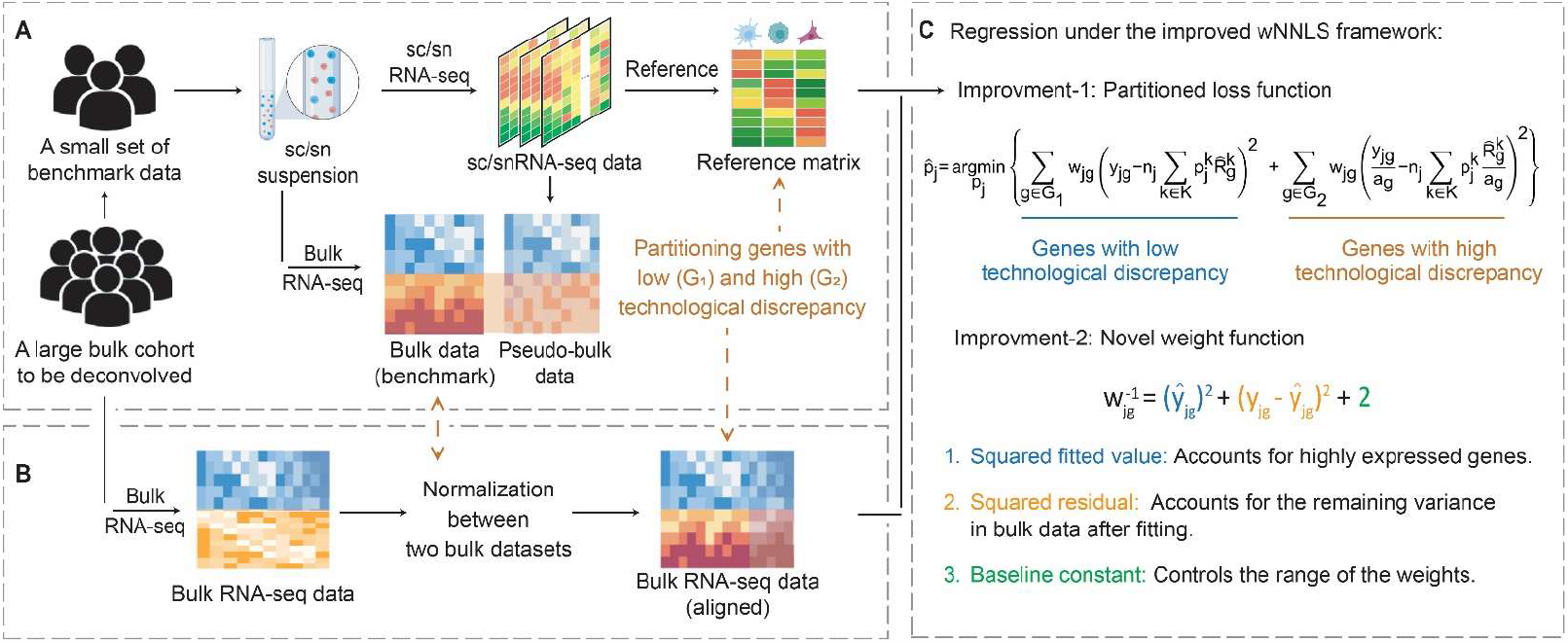
Overview of DeMixSC. The DeMixSC framework for deconvolution analysis of bulk RNA-seq data using sc/sn RNA-seq data as a reference. **A**, The framework starts with a benchmark dataset of matched bulk and sc/snRNA-seq data with the same cell-type proportions. Pseudo-bulk mixtures are generated from the sc/sn data. DeMixSC identifies DE and non-DE genes between the matched real-bulk and pseudo-bulk data. The non-DE genes are considered stably captured by both sequencing platforms (blue), while the DE genes are highly affected by technological discrepancy (orange). **B**, DeMixSC then employs a normalization procedure to perform the alignment between two bulk RNA-seq datasets (e.g., with ComBat). **C**, DeMixSC estimates cell-type proportions by regression under a weighted nonnegative least square (wNNLS) framework with two improvements: 1) partitioning and adjusting genes with high technological discrepancy, and 2) a new weight function. Here, *g* is the index of gene, *j* is the index of subject, *k* is the index of cell type, 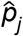 is the estimated cell-type proportions, *w*_*jg*_ is the weight, *n*_*j*_ is the normalization constant, 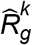 is the reference expression value derived from the sc/snRNA-seq data, *a*_*g*_ is the log_2_ transformed mean expression of the matched bulk and pseudo-bulk RNA-seq data, *y*_*jg*_ is the observed expression value in bulk RNA-seq data, and 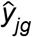 is the corresponding fitted value.

Here, *y*_*jg*_ is the observed expression value of gene *g* from subject *j* in bulk RNA-seq data, *n*_*j*_ is a normalization constant, 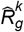 is the estimated cell-type-specific expression value of cell type *k* in the reference matrix derived from sc/snRNA-seq data, and *w*_*jg*_ is an associated weight. The gene sets *G*_*1*_ and *G*_*2*_ comprise genes with minimal and substantial impact by technological discrepancy, respectively. The first innovation of DeMixSC is the identification of and adjustment to the expression levels of genes that exhibit consistently high technological discrepancy (*G*_*2*_). We do so using a small representative benchmark dataset such as our special matched RNA-seq data from 24 retinal samples (Fig. 2A). DeMixSC uses a DE analysis between bulk and matched pseudo-bulk RNA-seq data to segregate genes with low inter-platform discrepancy (non-DE genes, *G*_*1*_) from those highly affected by technological discrepancy (DE genes, *G*_*2*_). It then employs a partitioned loss function and adjusts genes from *G*_*2*_ by rescaling their expressions by *a*_*g*_ (Figs. 2B, C) to mitigate the influence of technological discrepancy. Here we assign *a*_*g*_ as the log_2_-transformed mean expression of the matched bulk and pseudo-bulk RNA-seq data.

The second innovation of DeMixSC comes from our proposed function 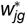, which is given by

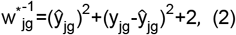

where 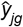 denotes the fitted expression value. This weight function comprises three terms: the squared fitted expression, the squared residual, and a baseline constant (Fig. 2C). The fitted part addresses genes with high expression levels. The squared residual accounts for the remaining variance after fitting. The baseline constant constrains the weight range. Previous weight functions contain either the first or the second part only^10,14–17^. We reason that both parts are needed to account comprehensively for variations across cells and samples. Hence, we introduce this new weight function. To demonstrate the advantages of the proposed weight based on the partitioned loss function, we compared the top 1,000 weighted genes from DeMixSC and MuSiC using the benchmark dataset. We observe that DeMixSC selects more genes with high technological discrepancy, while MuSiC selects slightly more genes with low discrepancy (Extended Data Figs. 2A, B). We calculated the between-cell-type collinearity among these genes to reflect the expression similarity of the reference profiles between different cell types (see Methods). The selected genes from DeMixSC present consistently lower between-cell-type collinearity than MuSiC across samples (Extended Data Figs. 2C, D, the median and median absolute deviation (MAD) of the reduction in Pearson correlation coefficients across 21 pairs of cell types are 0.37 and 0.11, Supplementary Table 3). This supports the ability of our proposed weight function in selecting genes that are informative to delineate cell-type-specific expressions, rather than those that minimize technological discrepancy but are less informative.

DeMixSC operates as a two-tier model in application. First, DeMixSC uses a specifically designed benchmark dataset to identify and adjust genes with high inter-platform discrepancy (Fig. 2A). Second, to deconvolve a large unmatched bulk RNA-seq dataset, DeMixSC aligns the large bulk cohort with the bulk RNA-seq data in the small benchmark dataset^33^ (Fig. 2B) to generalize the technological discrepancy detected and then runs the refined wNNLS framework for deconvolution (Fig. 2C). Our main prerequisite is a matched tissue type between the small benchmark dataset and the large targeted cohort.

### Comparing the estimation accuracy of DeMixSC with that of other existing deconvolution methods

Using our benchmark data, we compare the performance of DeMixSC with that of eight existing deconvolution methods^10–15,17,18^: AutoGeneS, BayesPrism, CIBERSORTx, DWLS, MuSiC, RNAseive, SCDC, and SQUID (see Methods, Fig. 3A). The retinal tissue samples in our benchmark dataset comprises ten distinct cell types. We focus our evaluation of different deconvolution methods on seven major cell types (Fig. 1A; amacrine cells, ACs; bipolar cells, BCs; Cone cells; horizontal cells, HCs; Müller glial cells, MGs; retina ganglion cells, RGCs; Rod cells), which on average account for 98% of the total cell population^34^.

**Figure 3.**
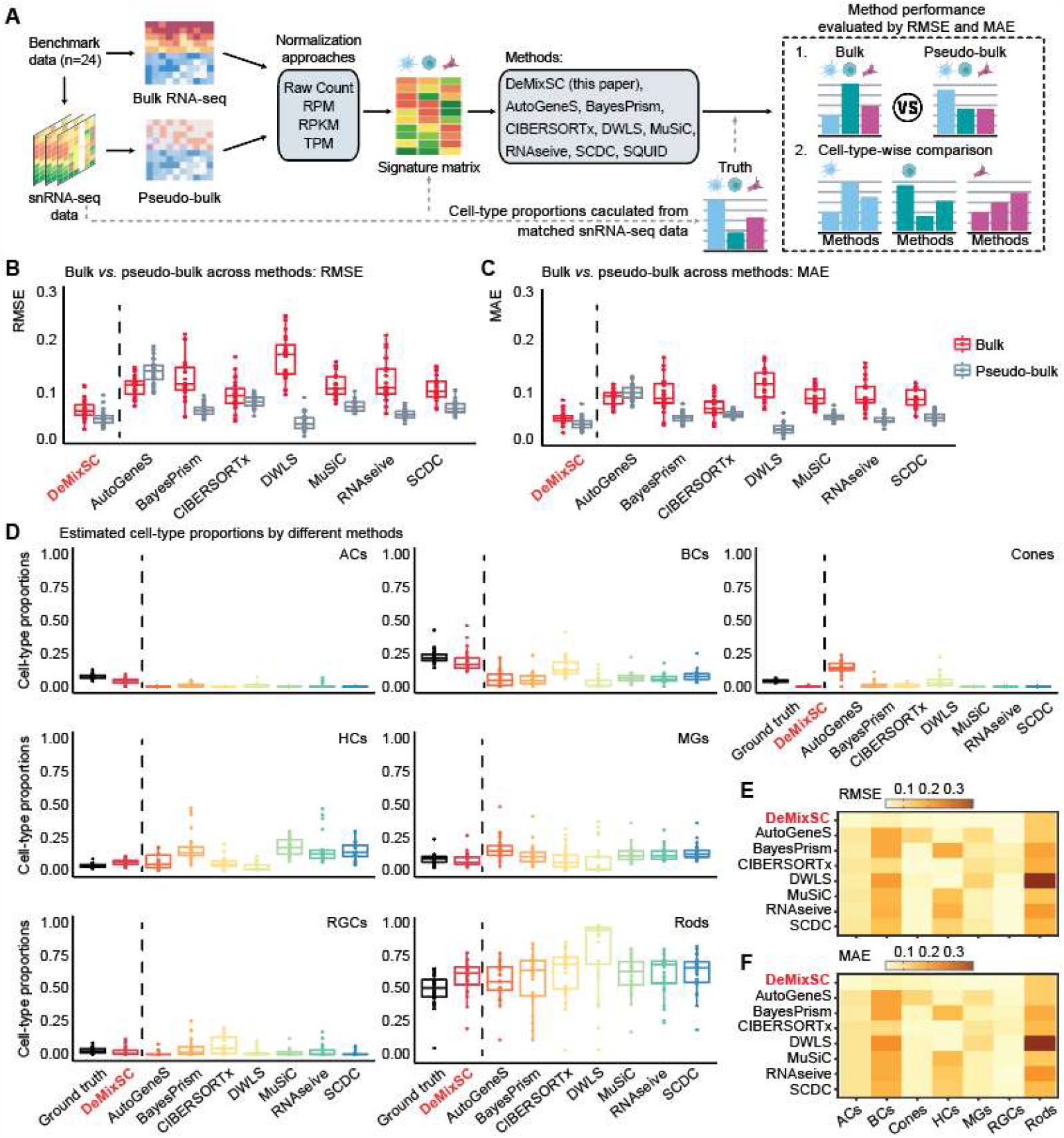
Compare the estimation accuracy of DeMixSC to existing deconvolution methods. **A**, Workflow for the deconvolution benchmarking design. We use benchmark data from retinal samples. The cell count proportions for each cell type are used as ground truth for the corresponding tissue samples. We assess the deconvolution performance of DeMixSC and seven existing methods for both bulk and pseudo-bulk mixtures. In addition to the raw counts, we also test RPM, RPKM, and TPM. The deconvolution performance is assessed by RMSE and MAE. **B** and **C**, Boxplots showing the deconvolution performance of eight deconvolution methods for the bulk and pseudo-bulk data. RMSE and MAE values are calculated across seven major cell types for each sample, with gray denoting pseudo-bulk and red denoting real-bulk. Smaller values indicate higher accuracy in proportion estimation. **D**, Boxplots showing the distributions of deconvolution estimates at the cell-type level for all 24 retinal samples. Each color corresponds to a given deconvolution method, with black denoting the ground truth, and each panel corresponds to a given cell type. **E** and **F**, An overview of deconvolution performance at the cell-type level across the eight methods using RMSE and MAE, respectively. Lighter colors correspond to lower RMSE or MAE values.

Overall, DeMixSC achieves the lowest root mean squared error (RMSE) and mean absolute error (MAE) scores in deconvolving bulk RNA-seq data, with mean values of 0.06 and 0.04 (Figs. 3B, C). Moreover, DeMixSC produces similar RMSE and MAE scores for deconvolving bulk and pseudo-bulk RNA-seq data (mean RMSE: bulk 0.06, pseudo-bulk 0.04; and mean MAE: bulk 0.04, pseudo-bulk 0.03). Those results suggest that DeMixSC adjusts well to undesired technological discrepancies. In contrast, existing methods perform poorly for bulk data but reasonably well for pseudo-bulk. In that sense, technological discrepancies that compromise deconvolution accuracy remain unaddressed by other existing approaches (Figs. 3B, C). Specifically, AutoGeneS shows higher RMSE and MAE scores for pseudo-bulk data, likely due to an inability to distinguish between Rod and Cone cells, which share largely similar expression profiles (Fig. 3D). DWLS excels in deconvolving pseudo-bulk samples but falls short for bulk RNA-seq data, possibly due to overfitting. Using the tree-based deconvolution in MuSiC or the ensemble option in SCDC does not improve their accuracy (Extended Data Fig. 3). CIBERSORTx presents overall reasonable performances in both bulk and pseudo-bulk data, likely because of its batch effect correction step. Looking further at the cell-type level, we observe systematic biases across methods. Most methods underestimate the proportions of ACs and BCs while overestimating HCs and MGs (Fig. 3D). DeMixSC accurately estimates the proportions of all seven major cell types and improves the deconvolution results for ACs, BCs, HCs, and MGs (Figs. 3D-F; RMSE: 0.04, 0.06, 0.03, 0.03 and MAE: 0.03, 0.05, 0.03, 0.02 for four cell types, respectively). Similar to most other methods, DeMixSC underestimates the proportion of Cone cells (true mean proportion at 0.04). In addition, we tested the robustness of these methods under varied data formats^35^, including RPM, RPKM, and TPM (see Methods), and found DeMixSC to be robust to data transformations (Extended Data Fig. 4). In line with previous benchmarking studies^29^, we find using raw counts as input is sufficient to obtain good results. Finally, SQUID delivered the least desirable results in this benchmarking study (mean RMSE and MAE in bulk data: 0.25 and 0.17). The issues with SQUID possibly lie in its data transformation step^28^, which has the potential to misrepresent gene expression profiles. In summary, our DeMixSC framework has achieved the most accurate deconvolution among the compared methods by successfully addressing the key issues of technological discrepancies between pairs of sequencing platforms.

### Applying DeMixSC to human peripheral retina bulk RNA-seq data

Age-related macular degeneration (AMD) is characterized by deterioration of retina and choroid that leads to substantial loss of visual acuity, with loss of Rod cells as a major manifestation. It is the leading cause of blindness among the elderly population globally^36^. However, the molecular and cellular events that underlie AMD remain poorly understood, impeding the development of effective treatments^37^. Understanding the molecular and cellular dynamics is essential for targeting the progression of AMD. We aim to examine cell-type proportion changes during AMD progression using bulk RNA-seq samples from 453 human peripheral retinas^7^ (see Methods). Among these retina samples, 105 have been scored in the Minnesota Grading System as grade 1 (MGS1), with 97 as MGS2, 88 as MGS3, and 61 as MGS4. An MGS1 rating indicates non-AMD healthy retina, and an MGS4 rating indicates AMD. MGS2 and MGS3 represent intermediate stages^38^.

We ran the two-tier DeMixSC to align the AMD cohort with the bulk data from our specialized benchmark dataset of retina samples, and then ran wNNLS to estimate cell-type proportions in the AMD cohort (see Methods, Fig. 4A). To achieve reliable deconvolution estimates, we constructed a consensus reference by integrating expression profiles from seven single-nucleus samples (see Methods). DeMixSC produces robust deconvolution estimates among the consensus and each individual single-nucleus reference at both cell-type and sample levels (Figs. 4B, C). The top-weighted genes identified by DeMixSC consistently maintain low between-cell-type collinearity across samples (see Methods, Extended Data Fig. 5). Consequently, DeMixSC achieves cell-type proportions (Fig. 4D) that are closer to experimental measures for non-AMD samples^39^, with an MAE of 0.03 (Supplementary Table 4). For comparison, we deconvolve the same cohort with MuSiC2^16^, CIBERSORTx^18^, and SQUID^10^, where MuSiC2 was chosen for its added ability to leverage conditionally stable genes from healthy references in analyzing diseased tissue. All three methods present biased estimates, especially in the proportion of Rod cells (Extended Data Fig. 6). DeMixSC identifies consistent changes in proportions in three major cell types over the axis of MGS1-4 (Extended Data Fig. 7), including a statistically significant decrease in the proportions of Rod cells (non-AMD: 0.57 vs. AMD: 0.53; t-test *P*-value = 0.02), as well as statistically significant increases in proportions of BCs (non-AMD: 0.16 vs. AMD: 0.17, t-test *P*-value = 0.05) and MGs (non-AMD: 0.12 vs. AMD: 0.15, t-test *P*-value < 0.001). It is known that the adult retina is not able to generate new cells^36,37,40^, so we hypothesize that it is the loss of Rod cells that results in a decreased total number of cells, hence increasing the Rod cell proportions as well as proportions of other cells in the AMD condition. Indeed, we find that losing 15% of all Rod cells can result in the observed subtle drop from 0.57 to 0.53 for the proportions of Rod cells, as well as the subtle increases in proportions of the other two cell types (see Methods).

**Figure 4.**
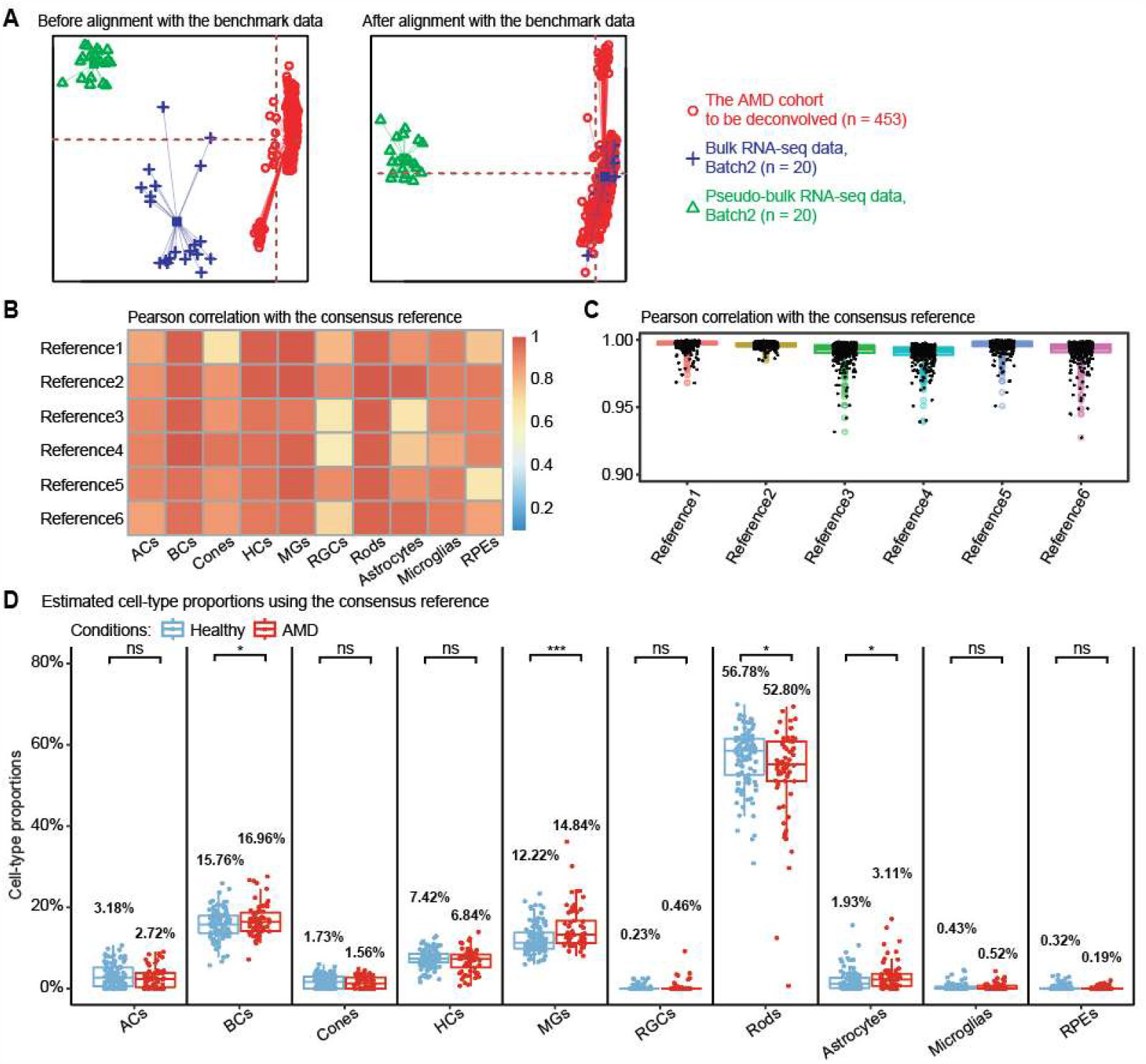
Using DeMixSC to deconvolve a large cohort of human peripheral retinal samples. **A**, PCA plots of both the retina cohort data and the benchmark data. Red denotes the bulk data to be deconvolved, blue denotes the benchmark bulk data, and green denotes the benchmark pseudo-bulk data. **B** and **C** demonstrate the robustness of DeMixSC to different reference matrices at both cell-type and sample levels. Higher correlation coefficients indicate better performance. **D**, Distributions of DeMixSC estimated cell-type proportions of *Ratnapriya et al*. data using consensus references. Each panel corresponds to a given cell type. The *P*-values for Student’s t-tests comparing the estimated cell-type proportions between non-AMD (healthy) and AMD groups are denoted as follows: not significant (ns), *P*-value >0.05; **\****P*-value ≤0.05; \*\**P*-value ≤0.01; and \*\*\**P*-value ≤0.001.

To demonstrate the biological significance of DeMixSC, we examined the DeMixSC-derived top-weighted gene for both non-AMD and AMD conditions (see Methods), revealing a substantial overlap of 93.8% in genes associated with DNA and RNA biosynthesis (Supplementary Table 5). Furthermore, DeMixSC adeptly pinpoints genes unique to each condition: cell adhesion genes in non-AMD^41^ and reactive oxygen species genes in AMD^42^ (Supplementary Table 5). These results suggest that DeMixSC not only identifies condition-stable genes but also delineates condition-specific signatures, enhancing deconvolution accuracy across conditions.

## Discussion

This study addresses the technological discrepancy between bulk and sc/sn RNA-seq data in order to improve the deconvolution accuracy of bulk RNA-seq data. We constructed a specialized benchmark dataset of healthy retina samples and thoroughly evaluated the impact of technological discrepancy on existing single-cell-based deconvolution methods^10–18^. Using this benchmark dataset, we introduce the DeMixSC deconvolution method that makes innovative improvements on the wNNLS framework to address the consistently observed technological discrepancy at the gene level. The distinct advantage of DeMixSC lies in its superior deconvolution accuracy and broad generalizability. As demonstrated in the benchmarking study, DeMixSC achieves more accurate estimates of cell-type proportions than other existing deconvolution methods. Importantly, DeMixSC is generalizable to deconvolving unmatched large bulk datasets, only requiring a small set of tissue-type-matched benchmark data. In our application to complex retina samples from patients with AMD, DeMixSC was able to accurately delineate 7 to 10 cell types and identified subtle yet critical changes in cell-type proportions of Rod, bipolar, and Müller glial cells. Our result highlights the utility of DeMixSC to capture cellular dynamics during disease progressions. DeMixSC is computationally efficient, completing the analysis of 453 AMD samples in under five minutes, and exhibits robust convergence against different starting values (see Methods, Extended Data Fig. 8). Our implementation of the DeMixSC framework is available at https://github.com/wwylab/DeMixSC.

Generation of the benchmark dataset in DeMixSC is crucial for accurate and reliable estimates of cell-type proportions. Our study employed a specifically tailored cDNA library preparation procedure to generate the benchmark dataset of retinal samples. A critical step in the data generation process is to ensure the ‘matchness’ of paired bulk and snRNA-seq data. In our procedure, the cDNA library for bulk RNA-seq was generated using the Smart-Seq v4 ultralow input RNA kit procedure, a protocol similar to that used in snRNA-seq. We advocate the preparation of ample tissue samples when executing this pipeline to ensure a reliable benchmark dataset.

The advance represented by DeMixSC is noteworthy, but there is potential room for improvements in future work. The key to DeMixSC rests on effectively identifying and down-weighting genes with high technological discrepancy. A potential challenge arises in gene identification when applying DeMixSC to tissue types (e.g., tumors) with high cellular plasticity. In that scenario, a stratified categorization of genes into three distinct groups can be beneficial: technologically stable genes, biologically stable genes (e.g., global tumor signature genes^6^), and the remaining unstable genes. Moreover, DeMixSC can be expected to gain from machine learning models to simultaneously identify and adjust genes. Additionally, alternative methods to ComBat^33^ for aligning the large cohort with the benchmark dataset can be considered when dealing with tumor samples, which often are highly heterogeneous with complex batch structures. Considering such potential adaptations, we anticipate that DeMixSC will prove useful in cancer research. By using a concise benchmark dataset derived from matched tissue specimens, DeMixSC can be leveraged to accurately deconvolve large bulk cohorts acquired through either surgical or biopsy samples. This capability can be expected to accelerate the discovery of cell subtypes and cell-type-specific markers among diverse patient groups with a variety of different types of cancer.

## Methods

### Human retina sample collection

These samples were obtained from 24 individuals between the ages of 73 and 91 who had passed away due to respiratory or heart failure or from a myocardial infarction (Supplementary Table 1). Human donor eyes were obtained through the Utah Lions Eye Bank. For this study, we included samples collected within six hours postmortem. Dissections of donor eyes were performed immediately following a previously published protocol^43^. Macular retinal tissue was collected using a six mm disposable biopsy punch (Integra, Cat # 33-37), flash-frozen, and stored at -80°C. Only one eye was used per donor, and donors with any history of retinal degeneration, diabetes, macular degeneration, or drusen were excluded from the study. Additionally, each donor underwent an ophthalmology check to ensure that the eye was in a healthy condition. Institutional approval for patients consent to donate their eyes was obtained from the University of Utah, and the study adhered to the principles of the Declaration of Helsinki. All retinal tissues were deidentified in accordance with HIPAA Privacy Rules.

### Generation of benchmark data from 24 human retinal samples

#### Single-nucleus mRNA sequencing

Nuclei were isolated with prechilled fresh-made RNase-free lysis buffer (10 mM Tris-HCl, 10 mM NaCl, 3 mM MgCl2, 0.02% NP40). The frozen tissue was resuspended and triturated to break the tissue structure in lysis buffer and homogenized with a Wheaton™ Dounce Tissue Grinder. Isolated nuclei were filtered with 40 μm Flow Cell Strainer and stained with DAPI (4′,6-diamidino-2-phenylindole, 10 μg/ml) before fluorescent cytometry sorting (FACS) on an FACSAria III Cell Sorter (BD, San Jose, CA, USA) in the Cytometry and Cell Sorting Core at Baylor College of Medicine (BCM). All single-nucleus RNA sequencing was performed at the Single Cell Genomics Core (SCGC) at BCM. Single-nucleus cDNA library preparation and sequencing were performed following the manufacturer’s protocols (https://www.10xgenomics.com). A single-nucleus suspension was loaded on a Chromium controller to obtain a single-cell GEMS (gel beads-in-emulsions) for the reaction. The snRNA-seq library was prepared with chromium single cell 3′ reagent kit v3 (10x Genomics). The product was then sequenced on an Illumina NovaSeq 6000 (https://www.illumina.com).

#### Bulk mRNA sequencing of retina single-nucleus suspension

To ensure the ‘matchness’ of paired bulk and snRNA-seq data, the mRNA library for bulk RNA-seq followed the same pipeline as for snRNA-seq. Specifically, matched samples with snRNA-seq were used for RNA isolation by applying TRIzol (Invitrogen) to the separated single-nucleus resuspension. cDNA was prepared from ∼1 ng of total RNA by using the Smartseq v4 ultralow input RNA Kit according to the manufacturer’s directions (Takara). The libraries were made using Nextera XT library prep (Illumina). Full-length RNA-seq was performed on NovaSeq 6000 sequencers according to the manufacturer’s directions (Illumina).

### Preprocessing of snRNA-seq and bulk RNA-seq data

Retina snRNA-seq UMI (unique molecular identifier) count matrices were obtained using CellRanger (version 3.1.0) following the official guide to estimate absolute counts and were then processed using the Seurat^44^ package (version 3.6.0). Specifically, for each snRNA-seq dataset, we first removed genes expressed in fewer than 5% of cells; then filtered out cells with either fewer than 500 total UMIs or 200 expressed genes, or more than 50% total UMI counts derived from mitochondrial genes. The total numbers of transcripts of each cell were then normalized to 10,000, followed by a natural log transformation. Highly variable genes were detected and used for principal component analysis (PCA). Cells were then clustered using the Seurat package at a resolution of 0.5.

For bulk RNA-seq data, the quality of raw sequencing data was first evaluated by FastQC (version 0.11.9), and low-quality reads and adapters were then trimmed by Trimmomatic (version 0.4.0). Next, reads that passed quality control were aligned to GRCh38 using the 2-pass mode of STAR (version 2.7.7b), and read counts were obtained by featureCount function from the Subread package (version 1.22.2) following the standard pipeline.

### Cell-type annotation for snRNA-seq data

Seven major cell types, including Cone cells, Rod cells, horizontal cells (HCs), bipolar cells (BCs), amacrine cells (ACs), retinal ganglion cells (RGCs), and Müller glia cells (MGs), were annotated using known marker genes^39,40^ (Extended Data Figs. 1A, B and Supplementary Table 2). For the deconvolution analysis of bulk AMD retinal samples^7^, we included additional 3 minor cell types, including astrocytes, microglia cells, and retina pigmental epithelium (RPE).

### Generation of ground truth proportion and pseudo-bulk mixtures

With each annotated snRNA-seq data, the true proportion of each cell type was calculated as the number of cells in the cell type divided by the total number of cells. Pseudo-bulk mixtures corresponding to each bulk were calculated by adding up the UMI counts from all the annotated cells per gene from the matched snRNA-seq data.

### Statistical analysis

We used paired Student’s t-tests to identify the differentially expressed (DE) genes between matched bulk and pseudo-bulk RNA-seq data. The *P*-values for DE analysis were adjusted for multiple testing by the Benjamini-Hochberg (BH) method^45^. We used Student’s t-tests to compare the estimated cell-type proportions between non-AMD and AMD conditions from different deconvolution methods. We used Wilcoxon rank-sum tests to compare the sequencing read depth between bulk and pseudo-bulk data. For all *P*-values in this study, significance levels were denoted as follows: no significant (ns), *P*-value >0.05; **\****P*-value ≤0.05; \*\**P*-value ≤0.01; and \*\*\**P*-value ≤0.001.

### DeMixSC deconvolution framework

DeMixSC is a reference-based model built upon the wNNLS deconvolution framework with several improvements. Our model explicitly requires a benchmark dataset for training. To begin with, we revisit the core equation of existing deconvolution methods^10,11,14–17,30^, which is

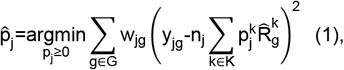

where *y*_*jg*_ is the observed expression value of gene *g* from subject *j* in the bulk RNA-seq data, 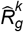is the estimated expression value of gene *g* and cell type *k* in the reference matrix derived from sc/snRNA-seq data, *w*_*jg*_ is the weight of each gene g from subject *j*, 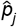 is the estimated vector of cell-type proportions, and *n*_*j*_ is a normalization constant. The main drawback of model (1) is that it does not address technological discrepancies observed in our benchmark data. To explain this, we split the squared term in Eq(1) into two components, and rewrite the model as

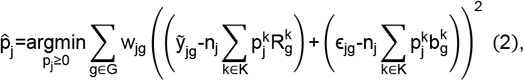

where 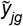 is the true expression value in the bulk data, 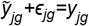, and 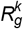 is the true cell-type-specific reference matrix, 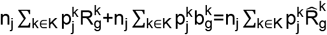, consists of the. The left component of Eq(2),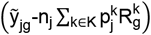, consists of the true bulk-level expression 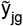 and the fitted value based on true cell-type-specific mean expression 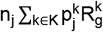. This component reflects the true estimation errors that we aim to minimize. The right component in Eq(2), 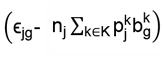, defines the difference in noise introduced by the bulk (*ϵ*_*jg*_) and the sc/snRNA-seq data (denoted by 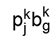 for cell type *k*). Therefore, the right component in Eq(2) represents the measurable technological discrepancy between sequencing platforms. Genes with higher levels of technological discrepancy contribute more to the right component (see Supplementary Note 1). Thus, when the technological discrepancy overtakes the true signal, instead of minimizing estimation errors (the left component), this model is geared towards minimizing the technological discrepancy (the right component) and is no longer fitting the expression profiles of individual bulk samples.

To address the issue of overcorrection for the technological discrepancy in model (1), we introduce DeMixSC, which estimates cell-type proportions by minimizing a partitioned loss function, as shown below:

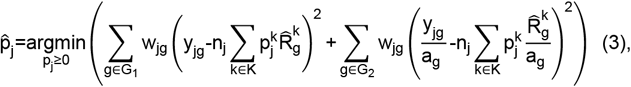

where G_1_ is a set of genes hardly affected by technological discrepancy, i.e., the non-DE genes between matched bulk and pseudo-bulk data, G_2_ contains genes highly affected by technological discrepancy, i.e., the DE genes between matched bulk and pseudo-bulk (see Supplementary Note 1), and a_g_ is the log_2_ transformed mean expression of the matched bulk and pseudo-bulk RNA-seq data. We use a_g_ to rescale the expression levels of DE genes to reduce the impact of the large differences in their error terms, as shown in Eq(2). Instead of directly filtering them out, our approach preserves those DE genes in our model, as they could still have biological significance and contribute to mixed expression levels. In addition, we introduce a new weight function 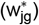 to reduce the influence of highly expressed genes and assign lower rankings for genes with large variances:

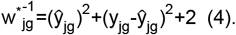

The current literature uses either the squared fitted value^10,15^ 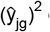 or the variance^14,16,17^ 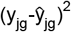 for weights, but never both. The constant 2 in Eq(4) is introduced for controlling the range of weights. Using the summation of the three terms as our new weight function improves model fit, accounts for variability, and enhances the statistical robustness of our framework. The detailed mathematical derivation is in Supplementary Note 1.

### Collinearity in top-weighted genes

To test the between-cell-type collinearity among samples, we selected the top 1,000 weighted genes from each deconvolved sample, and used corresponding expression levels to calculate the Pearson correlation coefficient between a pair of cell types and within the same sample.

### Convergence property of the DeMixSC algorithm

To evaluate how robust DeMixSC is against different initial values, we randomly selected a sample from the AMD cohort^7^ as a case study. To create different initial values, we set three different scale factors n = {100, 380, 1000}. For each scale factor, we chose 10 extreme starting values for the proportions 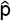, with the proportion of one out of 10 cell types being one and the rest being zero. Finally, we used the 30 values of 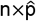 to initialize the wNNLS framework and then compared the estimates of DeMixSC.

### Data normalization of bulk mixtures

We applied the following data normalizations to the bulk raw count matrices^35^: (i) reads per million mapped reads (RPM); (ii) reads per kilobase of transcript, per million mapped reads (RPKM); and (iii) transcripts per million (TPM). Both RPKM and TPM include an additional step that uses the gene length to obtain normalized counts per million.

### Computational deconvolution

Nine deconvolution methods that use scRNA-seq as a reference were tested in our study^10–18^. We first used the default settings of each method as described in the GitHub repository or the websites (AutoGeneS: https://github.com/theislab/AutoGeneS, BayesPrism: https://github.com/Danko-Lab/BayesPrism, CIBERSORTx: https://cibersortx.stanford.edu/, DWLS: https://github.com/dtsoucas/DWLS, MuSiC: https://github.com/xuranw/MuSiC, MuSiC2: https://github.com/Jiaxin-Fan/MuSiC2/tree/master/R, RNAseive: https://github.com/songlab-cal/rna-sieve, SCDC: https://github.com/meichendong/SCDC, and SQUID: https://github.com/favilaco/deconv_matching_bulk_scnRNA). For CIBERSORTx, we followed the recommended built-in batch correction method for the deconvolution analysis of bulk samples (batch mode = S). Additionally, we evaluated the performance of the tree-guided deconvolution of MuSiC^17^ and the ensemble option of SCDC^14^. For tree-guided MuSiC, we first performed hierarchical clustering on the single-cell reference dataset; based on the hierarchical clustering results, we grouped Cone and Rod cells to form a mega cell cluster (Extended Data Fig. 3A), and each of the remaining cell types also formed a cluster. Cell-type-specific marker genes of cones and rods were obtained using FindAllMarkers function from Seurat^44^ package under the bimod likelihood ratio test. We ran MuSiC deconvolution first at the cell cluster level and then again within the Rod and Cone clusters. For the SCDC ensemble option, we ran deconvolution on SCDC with 3 different sc references; then, we ran the SCDC_ENSEMBLE function to obtain the ensemble deconvolution results.

### Evaluation metrics for the deconvolution performance

We evaluated the estimated cell proportions of each method using two metrics, root-mean-square error (RMSE) and mean absolute error (MAE), at both sample and cell-type levels: 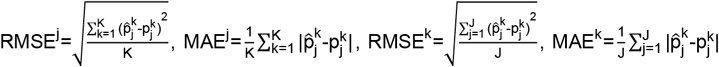, where 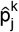 denotes the estimated cell proportion by the investigated method for cell type k and sample j, and 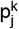 is the corresponding ground truth. We use J to denote the total number of samples and K to denote the total number of cell types. Smaller RMSE and MAE values indicate a better deconvolution performance.

### Quality control for the AMD cohort

The large AMD bulk cohort^7^ comprised 523 samples. We conducted quality control following the pipeline described in the original study. Briefly, samples were filtered out due to ambiguous clinical features (n=26), poor sequencing results (n=16), inconsistent genotyping results (n=14), and divergent ancestry (n=6). A total of 70 samples were removed, with a total of 453 samples remaining to be used to perform the deconvolution analysis.

### Consensus reference matrices for deconvolving the AMD cohort

For our deconvolution analysis of AMD retinal samples^7^, we generated a consensus single-nucleus reference by integrating seven samples from Batch-2 (Sample 5, 10, 12, 18, 19, 21 and 23, Supplementary Table 1). We selected these samples as they adequately represented these three minor cell types: astrocytes, microglia cells, and RPE. For each sample, we randomly selected up to 500 cells per cell type, using all available cells for types with fewer than 500 cells. Relative abundance and cell size for every cell type were calculated for each sample. A consensus reference matrix was subsequently derived by averaging relative abundance and cell size across the selected samples.

### Accounting for the total cell loss in the AMD cohort

The decline in overall cell count induced by Rod cell loss likely amplifies the cell-type proportions in the AMD samples^36,40,46^. To quantify the true proportion of Rod cell loss, we use x to represent the mean percentage of lost Rod cells in the AMD condition. With the mean estimated fraction of Rod cells is 0.57 in non-AMD and 0.53 in AMD (Fig. 4D), we derived the relation: 0.57(1-x)/(1-0.57x)=0.53. Solving for x gives 0.15. This 15% reduction in Rod cells in the peripheral AMD retina aligns well with biological evidence indicating the primary impact of AMD on the macular region^36^.

Then, we demonstrated the observed subtle increases in BCs and MGs are driven by the death of Rod cells. The mean estimated fractions of BCs and MGs in the non-AMD condition were 0.16 and 0.12 (Fig. 4D). Based on our Rod cell loss metric, the expected fractions in the AMD condition would be 0.16/(1-0.57*0.15)=0.17 for BCs and 0.12/(1-0.57x0.15)=0.13 for MGs, which were close to the DeMixSC’s estimates of BCs (0.17) and MGs (0.15) in the AMD condition (Fig. 4D).

### Gene ontology enrichment analysis of top-weighted genes

For each deconvolved sample from the non-AMD (n=105) and AMD (n=61) conditions, we identified the top 1,000 weighted genes. We then found the top 1,000 most frequently occurring genes across samples within each condition. To interpret the biological relevance of the gene list, we performed a Gene Ontology (GO) enrichment analysis using clusterProfiler4.0^47^ following the standard pipeline (http://yulab-smu.top/biomedical-knowledge-mining-book/GOSemSim.html).

## Data availability

The snRNA-seq data from the human retina tissue will be deposited and released at the Human Cell Atlas Data Portal. The raw data and count matrix of bulk RNA-seq data from human retinal tissue will be available at GEO under accession number: GSE175937.

## Code availability

DeMixSC is freely available as an R package and can be downloaded from our GitHub repository: https://github.com/wwylab/DeMixSC. A tutorial for DeMixSC is available at https://wwylab.github.io/DeMixSC/.

## Declarations

### Ethics approval and consent to participate

Institutional approval for patient consent to donate their eyes was obtained from the University of Utah, and the study adhered to the principles of the Declaration of Helsinki. All retinal tissues were deidentified in accordance with HIPAA Privacy Rules.

### Consent for publication

All authors have approved the manuscript for submission.

## Acknowledgements

S.G. is supported by Human Cell Atlas Seed Network - Breast, Chan Zuckerberg Institute, MD Anderson Colorectal Cancer Moon Shot Program, DoD PC210079. X.L. is supported by R01CA239342. A.K. is supported by 5T32CA096520-15. J.P.S. is supported by the Cancer Prevention and Research Institute of Texas as a CPRIT Scholar in Cancer Research and by National Institutes of Health (K22CA234406). R.C. is supported by CZI (CZF2019-02425), National Eye Institute (R01EY022356, R01EY018571), and Retinal Research Foundation. W.W. is supported by Human Cell Atlas Seed Network - Retina, Chan Zuckerberg Institute, NIH R01CA268380, DoD PC210079, P30CA016672. This work is also supported by CPRIT Single Core grant RP180684 and the Cytometry and Cell Sorting Core at Baylor College of Medicine with funding from the CPRIT Core Facility Support Award (CPRIT-RP180672), the NIH (CA125123 and RR024574) and the assistance of Joel M. Sederstrom. Single-nucleus RNA sequencing was performed at the Single Cell Genomics Core at BCM partially supported by NIH shared instrument grants (S10OD018033, S10OD023469, S10OD025240), P30CA125123, P30EY002520, and CPRIT Comprehensive Cancer Epigenomics Core Facility RP200504.

## Authors’ contributions

W.W. and R.C. supervised the research. W.W. and R.C. conceived the ideas and designed the study. S.G. and X.L. developed the method, implemented the R package, and conducted all the analysis. X.C. conducted the sequencing experiments and generated the paired bulk and single-nuclei RNA-seq data for benchmarking purposes. Y.J. and S.J. help with the initial development of the method and evaluation. Q.L. and Y.L. help with the sequencing experiments. L.A.O., I.K.K., A.A., S.K., J.P.S., M.M.D., and R.C. provided samples, advised on the study design, and assisted with the interpretation of results. A.K. performed the initial benchmarking analysis. J.N.W. and R.C. contributed technical suggestions. W.W., S.G., and X.L. wrote the paper, with input from all authors. All authors reviewed and approved the final version of the manuscript.

---

**Extended Data Figure 1.**
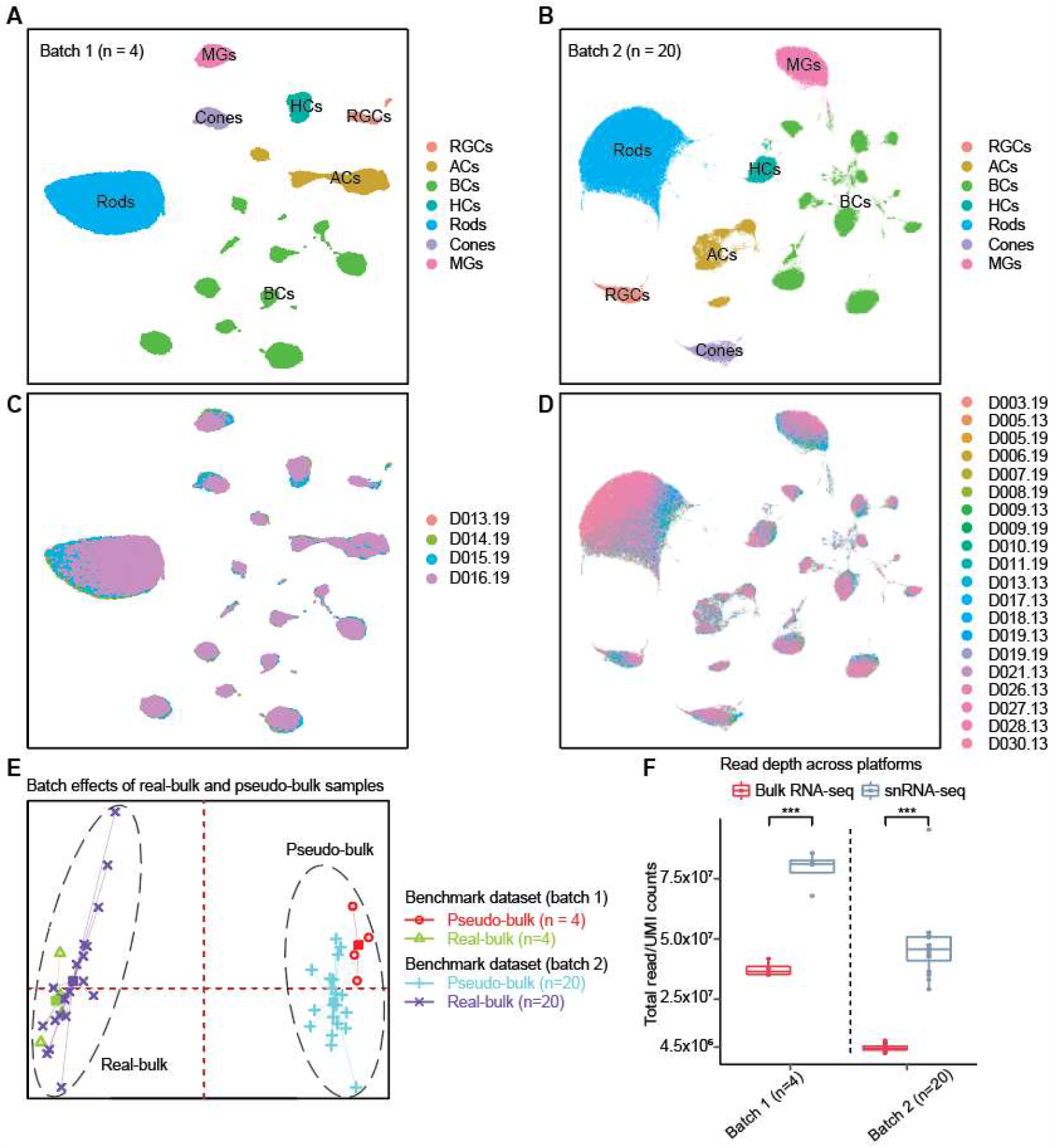
Overview of the matched bulk and snRNA-seq data. **A** and **B**, UMAP projection of snRNA-seq data from 4 healthy retinal samples in batch-1 and 20 healthy retinal samples in batch-2, annotated by cell types. **C** and **D**, UMAP projection of snRNA-seq data from 4 healthy retinal samples in batch-1 and 20 healthy retinal samples in batch-2, annotated by sample IDs. Cells were clustered by their biological annotations instead of sample origins, suggesting negligible batch effects. **E**, Distribution of the first two principal components for the matched real-bulk and pseudo-bulk RNA-seq data in the benchmark dataset. **F**, Boxplot showing the raw read depth between bulk and pseudo-bulk RNA-seq data from batch-1 and batch-2. The *P*-values for Wilcoxon rank-sum tests comparing sequencing read depth between bulk and pseudo-bulk data are denoted as follows: **\****P*-value ≤0.05; \*\**P*-value ≤0.01; and \*\*\**P*-value ≤0.001.

**Extended Data Figure 2.**
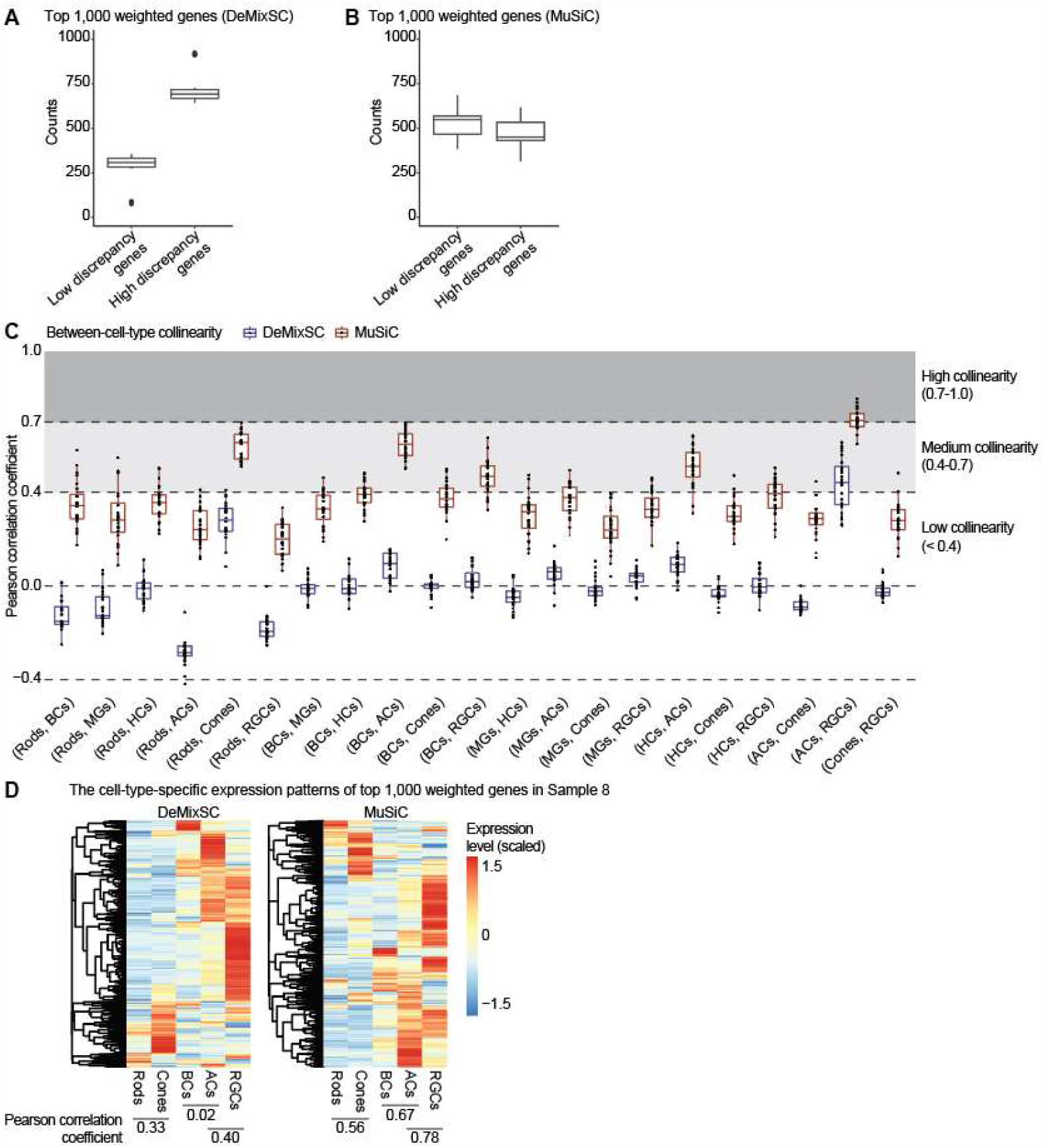
DeMixSC maintains low collinearity among top-weighted genes in the benchmark dataset. **A** and **B**, Boxplots showing the numbers of low and high discrepancy genes within the top 1,000 weighted genes across samples: DeMixSC **(A)** and MuSiC **(B). C**, Boxplot showing the between-cell-type collinearity among 21 cell-type pairs in the benchmark dataset across 24 retinal samples, as measured by Pearson correlation coefficient. The cell-type pairs are ordered according to their cell-type proportions within the tissue sample. MuSiC (red boxes) exhibits overall higher collinearity, especially for (Rods, Cones), (ACs, BCs), and (ACs, RGCs). In contrast, DeMixSC (blue boxes) presents only a slightly elevated correlation for (ACs, RGCs). The degree of collinearity is categorized as high (≥0.7), medium (0.4-0.7), and low (<0.4). **D**, Heatmaps showing cell-type-specific expression patterns of the top 1,000 weighted genes, using Sample 8 as an example for a mid-level correlation.

**Extended Data Figure 3.**
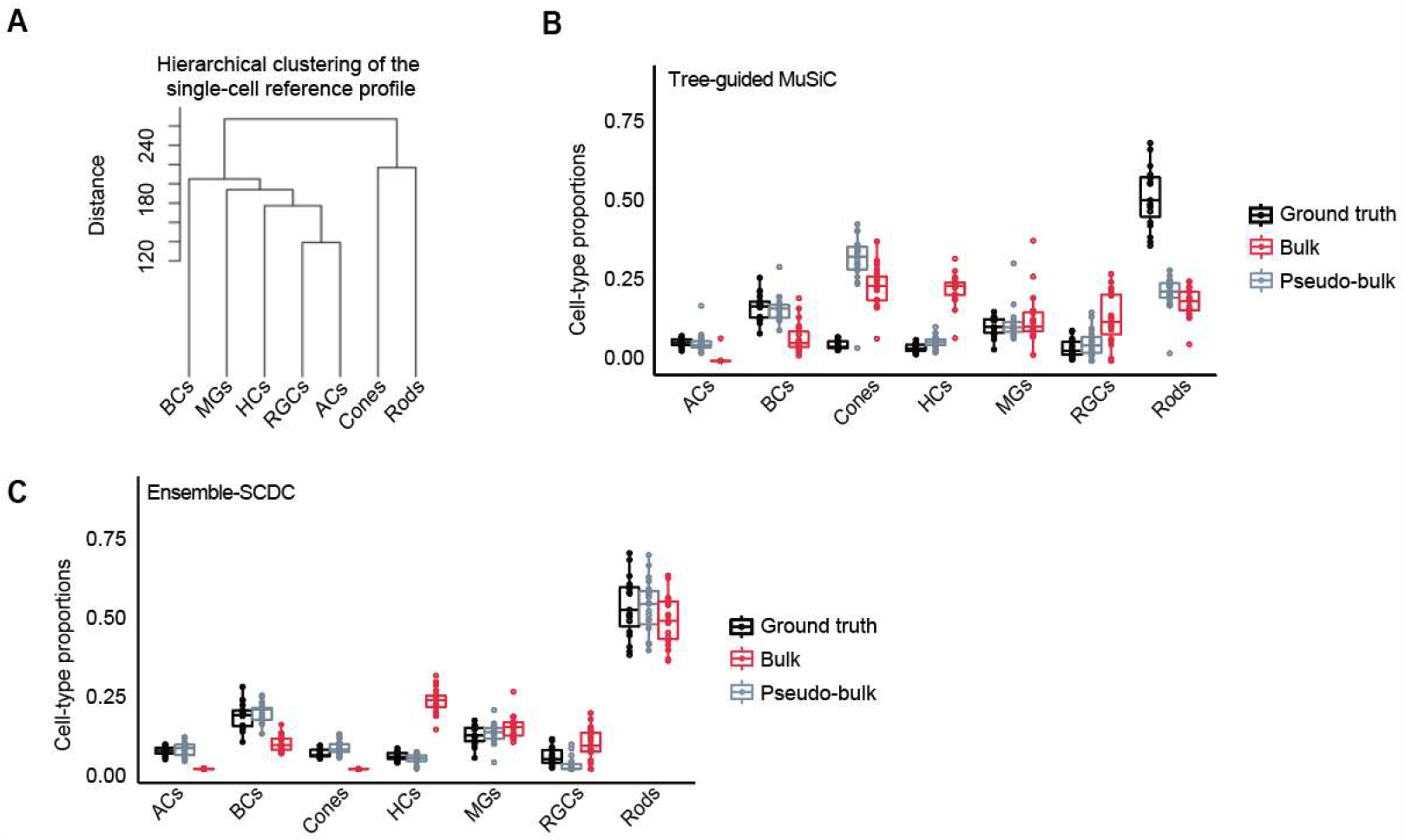
Deconvolution performance of the tree-guided MuSiC and Ensemble-SCDC. **A**, Hierarchical clustering of the cell-type-specific reference matrix. **B**, Boxplot showing the distributions of estimated cell-type proportions from the benchmark data using the tree-guided MuSiC. **C**, Boxplot showing the distributions of estimated cell-type proportions from the benchmark data using the SCDC ensemble mode. Black denotes the ground truth estimated using the snRNA-seq data. Gray denotes estimates from the pseudo-bulk RNA-seq data, and red denotes estimates from the matched bulk RNA-seq data.

**Extended Data Figure 4.**
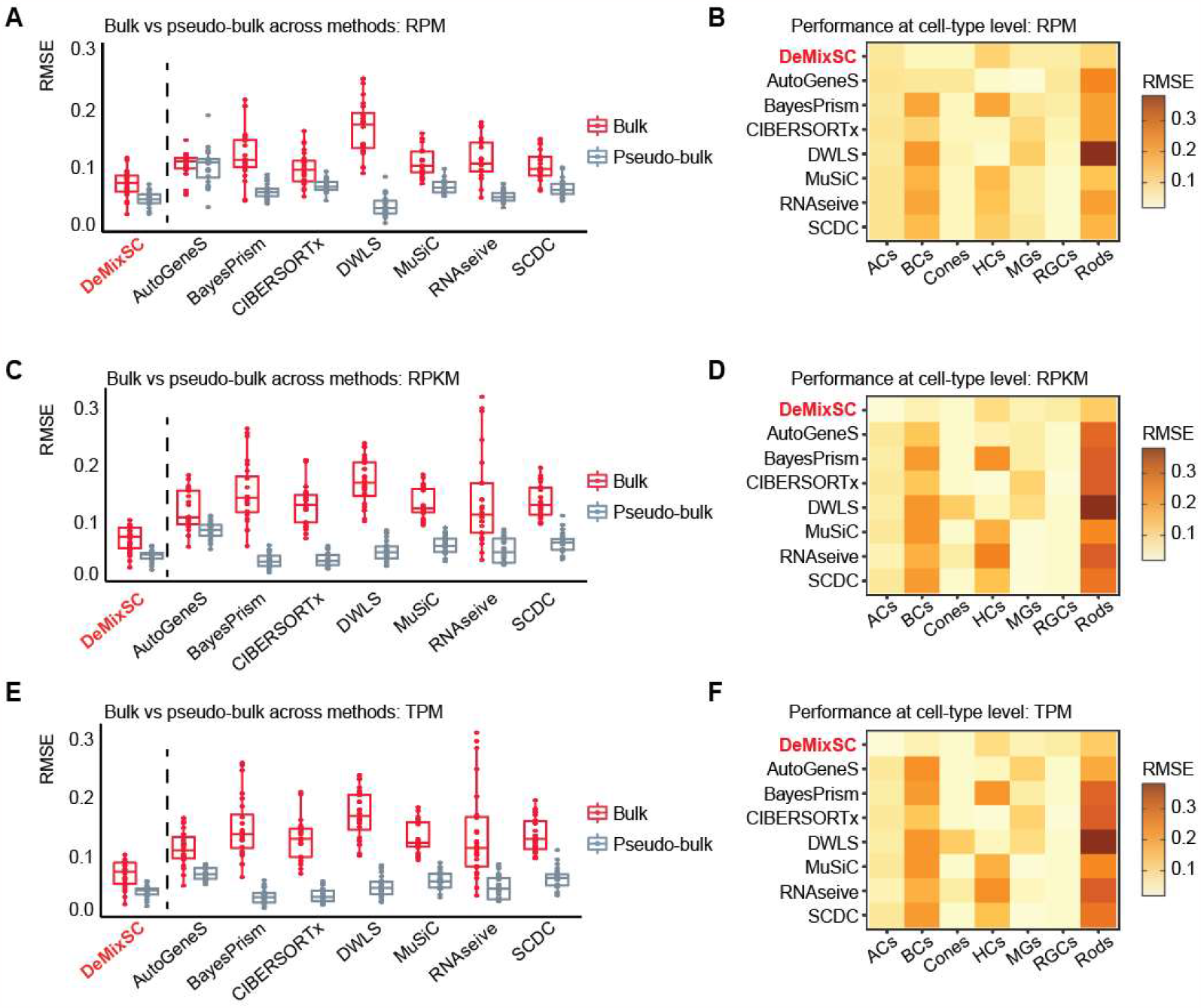
Impact of data normalization on the deconvolution performance. **A, C**, and **E**, Boxplots showing the deconvolution performance across DeMixSC and seven current single-cell-based deconvolution methods for bulk and pseudo-bulk mixtures. RMSE values are calculated across seven major cell types for each sample, with gray denoting pseudo-bulk and red denoting real-bulk. Smaller values indicate higher accuracy in proportion estimation. **B, D**, and **F**, Heatmaps showing the deconvolution performance at the cell-type level across the eight methods using RMSE. Lighter colors correspond to lower RMSE values. Each panel corresponds to a normalization strategy: RPM (**B**), RPKM (**D**), and TPM (**F**).

**Extended Data Figure 5.**
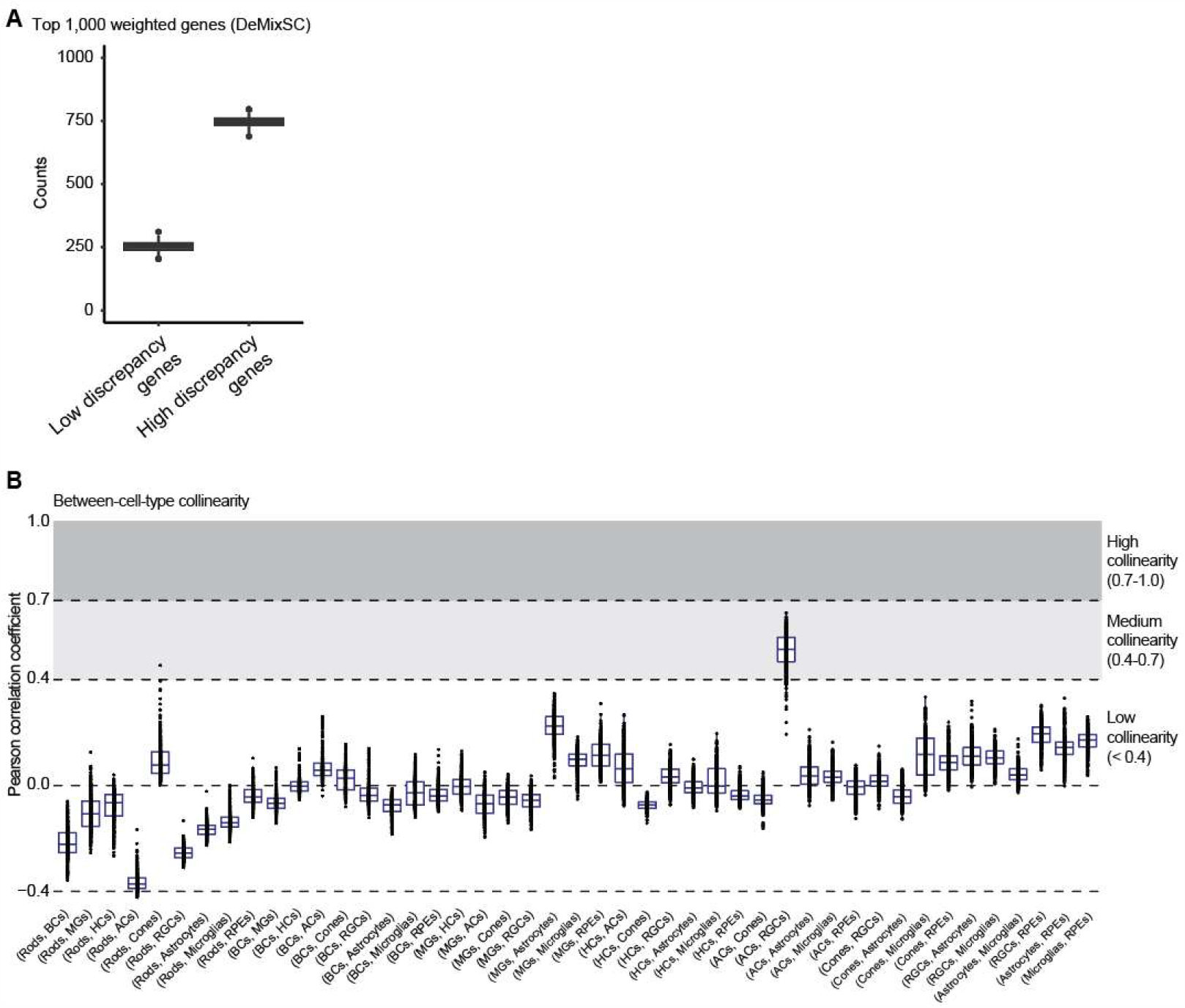
DeMixSC maintains low collinearity among top-weighted genes in the AMD cohort. **A**, Boxplot showing numbers of low and high discrepancy genes within the top 1,000 weighted genes for each sample. **B**, Boxplot showing the between-cell-type collinearity among 45 cell-type pairs (for 10 cell types) in the AMD cohort across 453 samples, as measured by Pearson correlation coefficient. The cell-type pairs are ordered according to their cell-type proportion within the tissue sample. Most of the cell-type pairs exhibit low collinearity, especially for pairs of major cell types on the left of the plot. Only the correlations between ACs and RGCs are slightly higher. A similar observation is made when deconvolving the benchmark data. The degree of collinearity is categorized as high (≥0.7), medium (0.4-0.7), and low (<0.4).

**Extended Data Figure 6.**
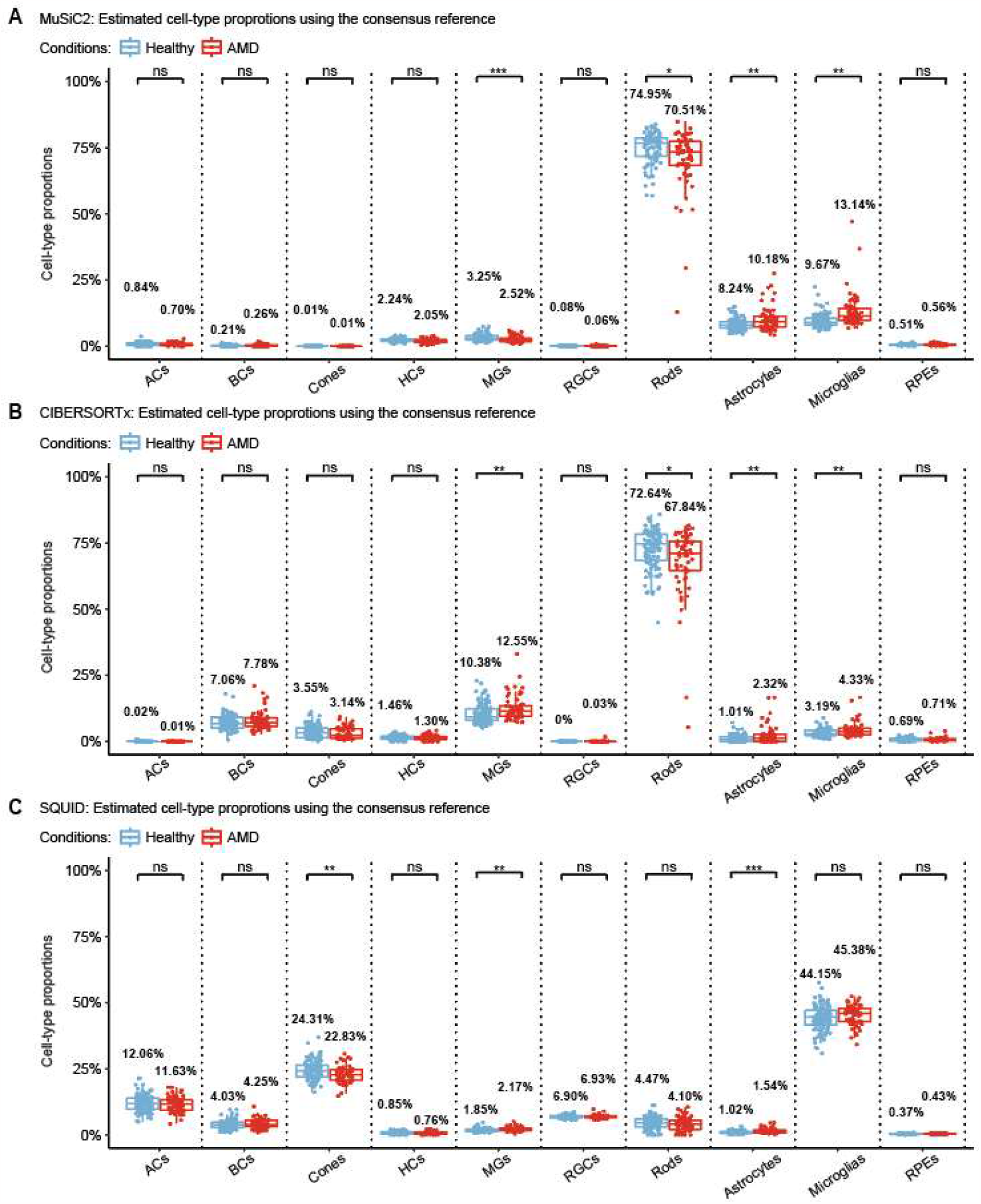
Cell-type proportion estimates for the AMD cohort with existing methods. **A, B**, and **C**, Boxplots showing the distributions of cell-type proportion estimates for non-AMD retina vs. AMD retina from MuSiC2 (**A**), CIBERSORTx (**B**), SQUID (**C**). The *P*-values for Student’s t-tests comparing the estimated cell-type proportions between non-AMD (healthy) and AMD groups are denoted as follows: not significant (ns), *P*-value >0.05; **\****P*-value ≤0.05; \*\**P*-value ≤0.01; and \*\*\**P*-value ≤0.001.

**Extended Data Figure 7.**
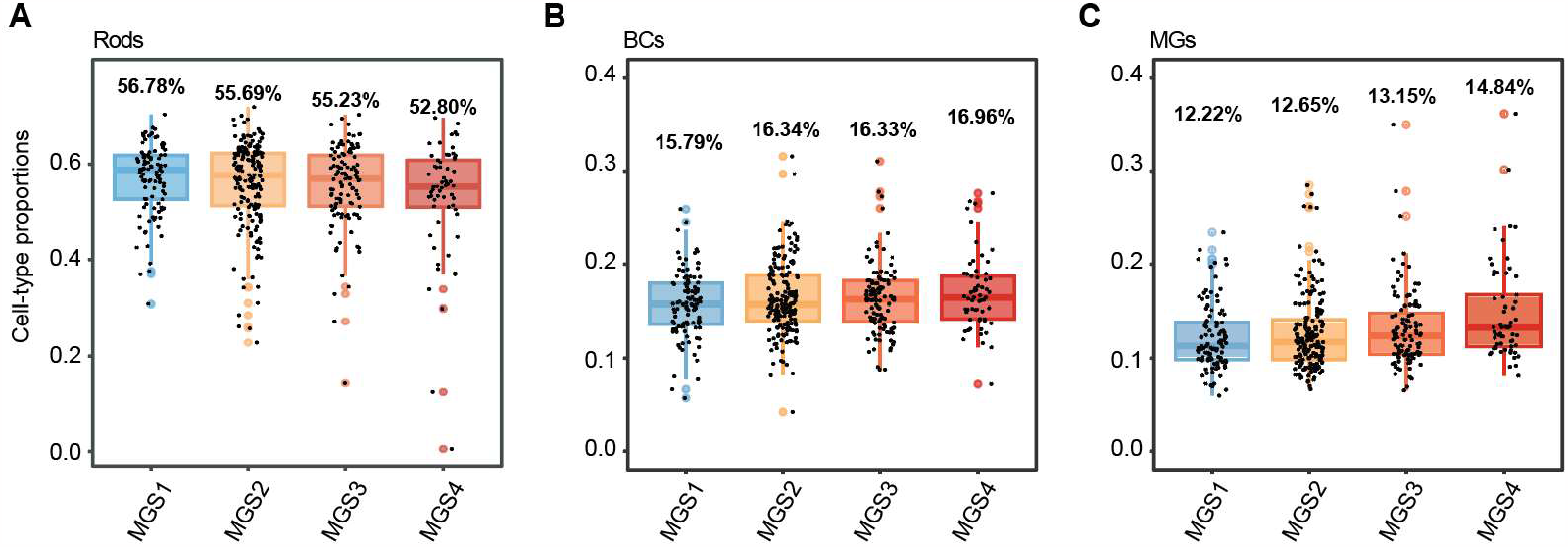
DeMixSC recovers a dynamic shift in cell-type proportions during the AMD progression. **A, B**, and **C**, Boxplots showing the distributions of cell-type proportion estimates across different MGS stages from MGS1 to MGS4. Each panel corresponds to a given cell type: Rods (**A**), BCs (**B**), and MGs (**C**).

**Extended Data Figure 8.**
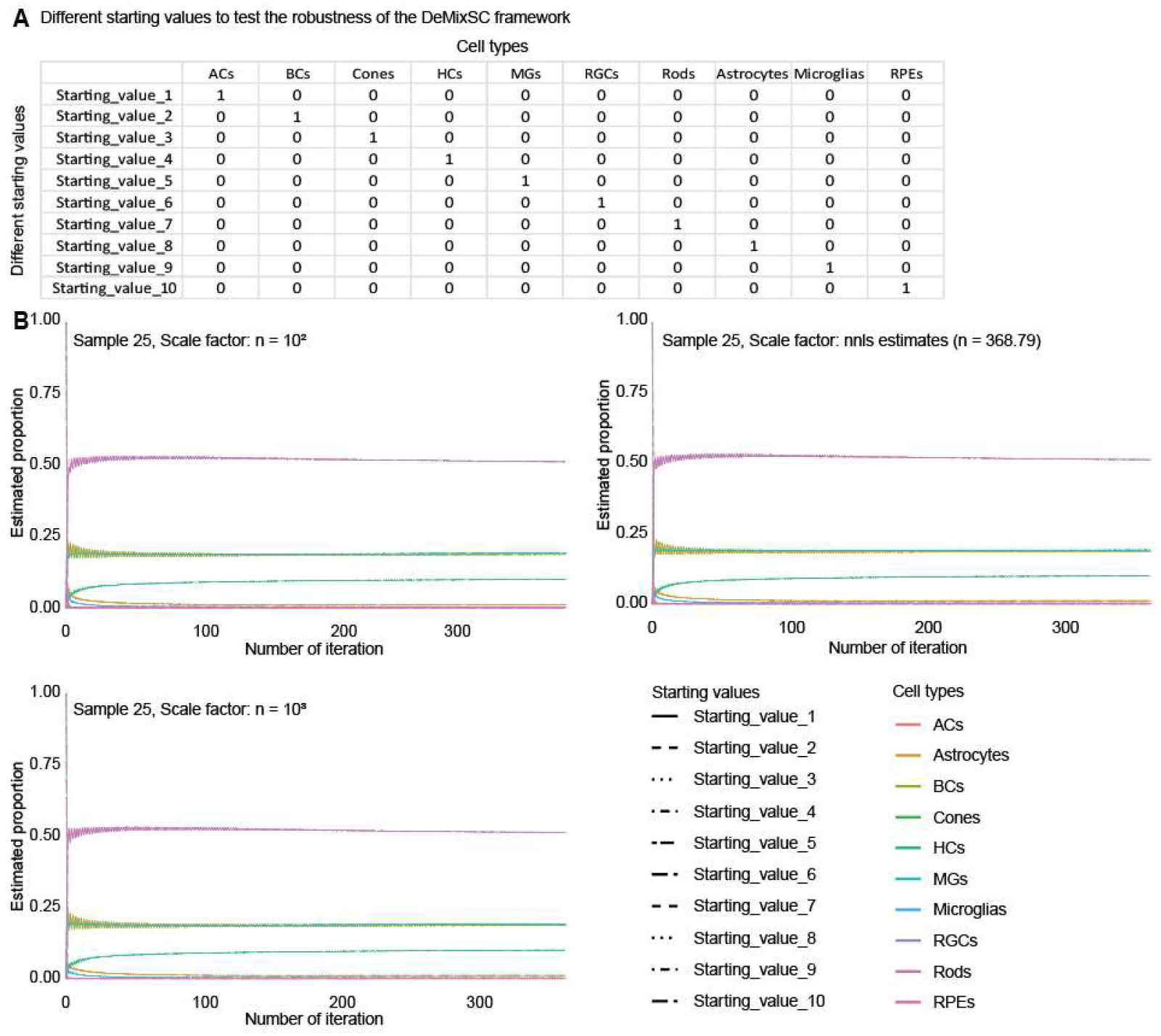
Convergence of DeMixSC with different starting values. **A**, A list of different starting values across ten cell types. **B**, Trace plots of estimated proportions over iterations.

